# Trefoil factors share a lectin activity that defines their role in mucus

**DOI:** 10.1101/859785

**Authors:** Michael Järvå, James P. Lingford, Alan John, Nichollas E. Scott, Ethan D. Goddard-Borger

## Abstract

The trefoil factors (TFFs) are disulfide-rich mucosal peptides that protect the epithelium by promoting cell migration and increasing the viscoelasticity of the mucosa. Here we show that all TFFs are divalent lectins that recognise the GlcNAc-α-1,4-Gal disaccharide, which terminates type-III mucin-like O-glycans. Structural, mutagenic and biophysical data support a model of mucus viscoelasticity that features non-covalent cross-linking of glycoproteins by TFFs.

---

The three human TFFs (**Fig. 1a**) (TFF1, TFF2 and TFF3)^1^ are ubiquitous in mucosal environments. They share substantial sequence similarity, suggestive of a conserved function, yet their biological roles are not redundant^2^. They protect the mucosal epithelium by increasing mucus viscoelasticity^3,4^ and enhancing epithelial restitution^5–9^. TFF overexpression is a hallmark of chronic inflammatory diseases of the respiratory tract^10–12^. The contribution of TFFs to the progression of these diseases is unclear, though any increase in mucus viscoelasticity might be expected to facilitate the formation of mucus plug obstructions in the airways. Exploring the role of TFFs in these and other biological contexts is confounded by an incomplete understanding of how they modulate the physical properties of mucus and a dearth of tools for inhibiting TFF activity^2^.

**Figure 1.**
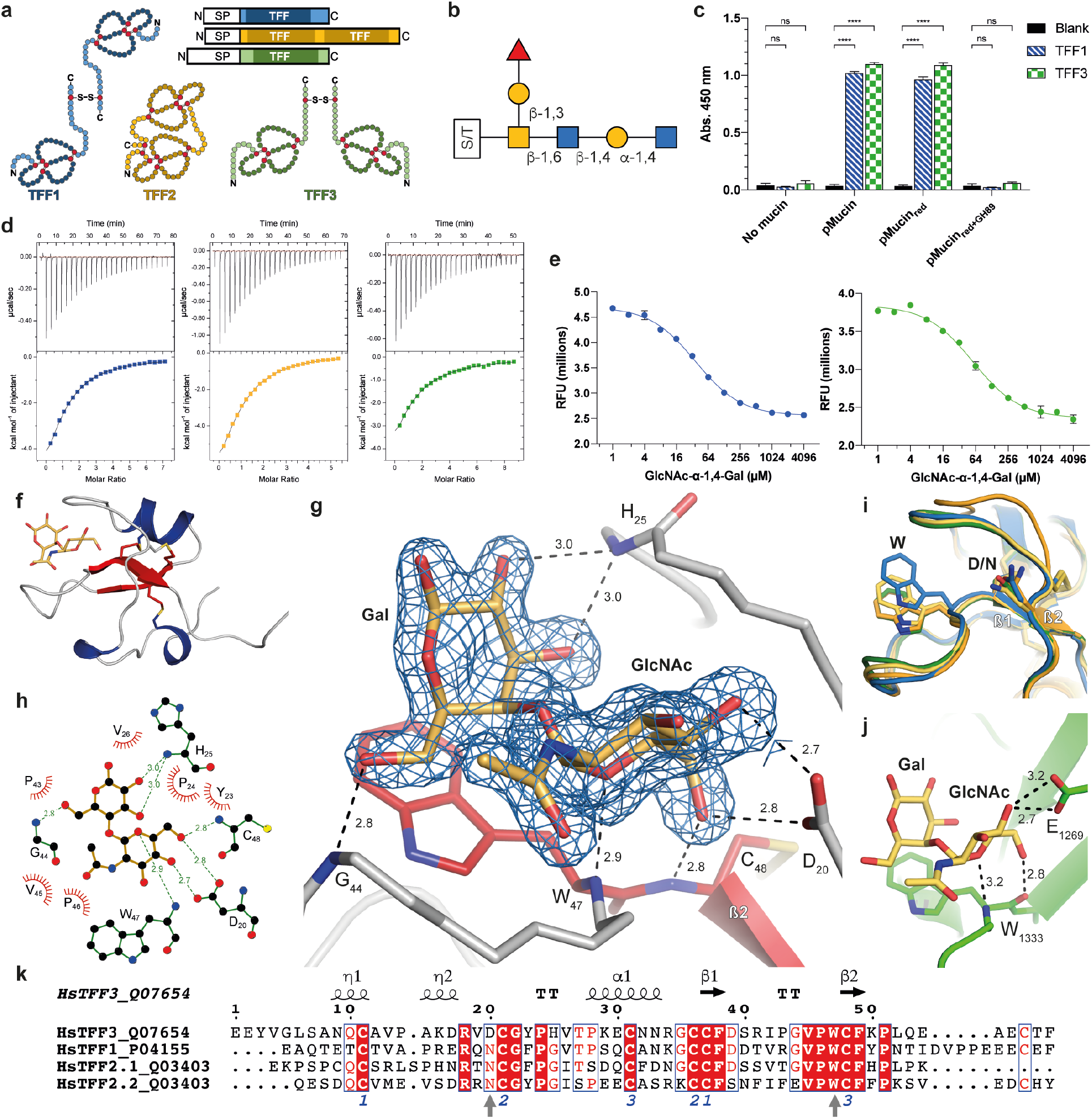
(a) Schematics and protein topology of human TFF1-3. (b) The α-GlcNAc-capped Core 2 O-glycan identified as a ligand for TFF2^14^. (c) ELISA data demonstrating the ability of mTFF1bio and mTFF3bio to bind to pMucin and pMucinred but not pMucinred+GH89. Error bars indicate standard deviation of three replicates. ns = p>0.05, **** = p<0.0001 (d) Representative ITC isotherms of mTFF1 (blue), TFF2 (orange) and mTFF3 (green) titrated with GlcNAc-α-1,4-Gal. (e) mTFF1 and mTFF3 binding to GlcNAc-α-1,4-Gal as determined by a tryptophan fluorescence quenching assays. Error bars indicate standard deviation of three replicates. (f) Structure of TFF3–GlcNAc-α-1,4-Gal. (g) Interactions between GlcNAc-α-1,4-Gal and TFF3 (distances in Å) and Fo-Fc omit map contoured at 1.5σ around the ligand. (h) Schematic representation of the interactions between TFF3 and GlcNAc-α-1,4-Gal. (i) Superposition of TFF3–GlcNAc-α-1,4-Gal (green), TFF1 (blue) and the trefoil domains from porcine TFF2 (orange and yellow) (PDBID: 2PSP). The glycan binding side chains are shown as sticks. (j) A CBM32 domain (PDBID: 4A6O) in complex with GlcNAc-α-1,4-Gal and their interactions (distances in Å). (k) Sequence alignment of the trefoil domains in human TFFs. Numbers indicate disulfide connectivity, and vertical arrows indicate conserved ligand-binding residues.

An extensive list of TFF binding partners has accumulated within the literature and includes many cell surface receptors and mucosal glycoproteins^2,13^. In the context of the mucus-thickening properties of the TFFs, the soluble mucins are the most relevant binding partners. This is complemented by the observation that TFF2 interacts with the α-D-GlcNAc-terminated O-glycans of mucins **(Fig. 1b)**^14^. It remains unknown what the simplest glycan structure required for TFF2 binding is, how tightly it binds, if TFF1 and TFF3 share this property and, if so, how the different biological activities of these closely-related peptides arise.

To establish that TFF1 and TFF3 possessed the same α-GlcNAc-dependent binding activity as TFF2, we started by preparing site-selectively biotinylated monomeric TFF1 and TFF3 (mTFF1_bio_ and mTFF3_bio_) (**Supplementary Table 1**) and three mucin preparations: commercially-available type-III porcine gastric mucin (pMucin), reduced and alkylated pMucin (pMucinred), and pMucin_red_ digested with an α-D-N-acetyl-glucosaminidase from *Clostridum perfringens* (*Cp*GH89) to remove terminating α-GlcNAc (pMucin_red+GH89_). ELISA performed using these reagents **(Fig. 1c)** detected robust binding of mTFF1_bio_ and mTFF3_bio_ to both pMucin and pMucin_red_, while no binding was observed for pMucinred+GH89. This established that all TFF–mucin interactions are independent of mucin protein structure but do require α-GlcNAc.

We next sought to determine the minimal carbohydrate structure required for TFF binding and the affinity of this interaction for each TFF. Isothermal calorimetry (ITC) demonstrated that the simple monosaccharides D-GlcNAc, benzyl α-D-GlcNAc and 4-nitrophenyl α-D-GlcNAc had no detectable affinity for TFF1-3. The larger GlcNAc-α-1,4-Gal disaccharide, which is the terminating structure unique to type-III mucins, bound to mTFF1, TFF2 and mTFF3 with a K_d_ of 49 ± 4 μM, 44 ± 8 μM and 65 ± 5 μM, respectively **(Fig. 1d, Supplementary Table 2 and Supplementary Fig. 1)**. Other α-1,4-linked disaccharides, such as maltose (Glc-α-1,4-Glc) and Gal-α-1,4-Gal, failed to show any affinity for the TFFs by ITC. These results were corroborated by a tryptophan fluorescence quenching assay, which gave K_d_ values for mTFF1 and mTFF3 of 37 ± 2 μM, and 58 ± 1 μM, respectively **(Fig. 1e)**.

Initial attempts to co-crystallise TFFs and their shared ligand (GlcNAc-α-1,4-Gal) were confounded by the exceptional solubility of these proteins. Eventually we obtained mTFF1 crystals that only yielded an apo structure (**Supplementary Table 3 and Supplementary Fig. 2**). Following mTFF3 surface lysine methylation, a crystal of the mTFF3–GlcNAc-α-1,4-Gal complex was obtained and the structure determined to a resolution of 1.55Å **(Fig. 1f, Supplementary Table 3)**. The disaccharide is accommodated within a hydrophobic cleft of TFF3 with ligand binding driven by sterical fitting and hydrogen bonds to the peptide backbone **(Fig. 1g,h)**. Only two side chains, Asp20 and Trp47, make direct contacts with the disaccharide: Asp20 makes a bidentate hydrogen-bonding interaction between O-4 and O-6 of the non-reducing α-GlcNAc, while C-H–π interactions are made between the indole ring of Trp47 and the reducing Gal. Trp47 undergoes considerable motion between liganded and unliganded forms **(Fig. 1i)**. These two residues are conserved in TFF1-3: though TFF1 and TFF2 feature an Asn in place of Asp20 **(Fig. 2k)**. This binding mode is remarkably similar to that of the bacterial carbohydrate-binding module (CBM) family 32 protein that binds the GlcNAc-α-1,4-Gal glycan (Kd of 72 μM) in the same conformational pose, despite sharing no sequence, structural or ancestral commonalities with the TFFs (**Fig. 2j**)^15^.

**Figure 2.**
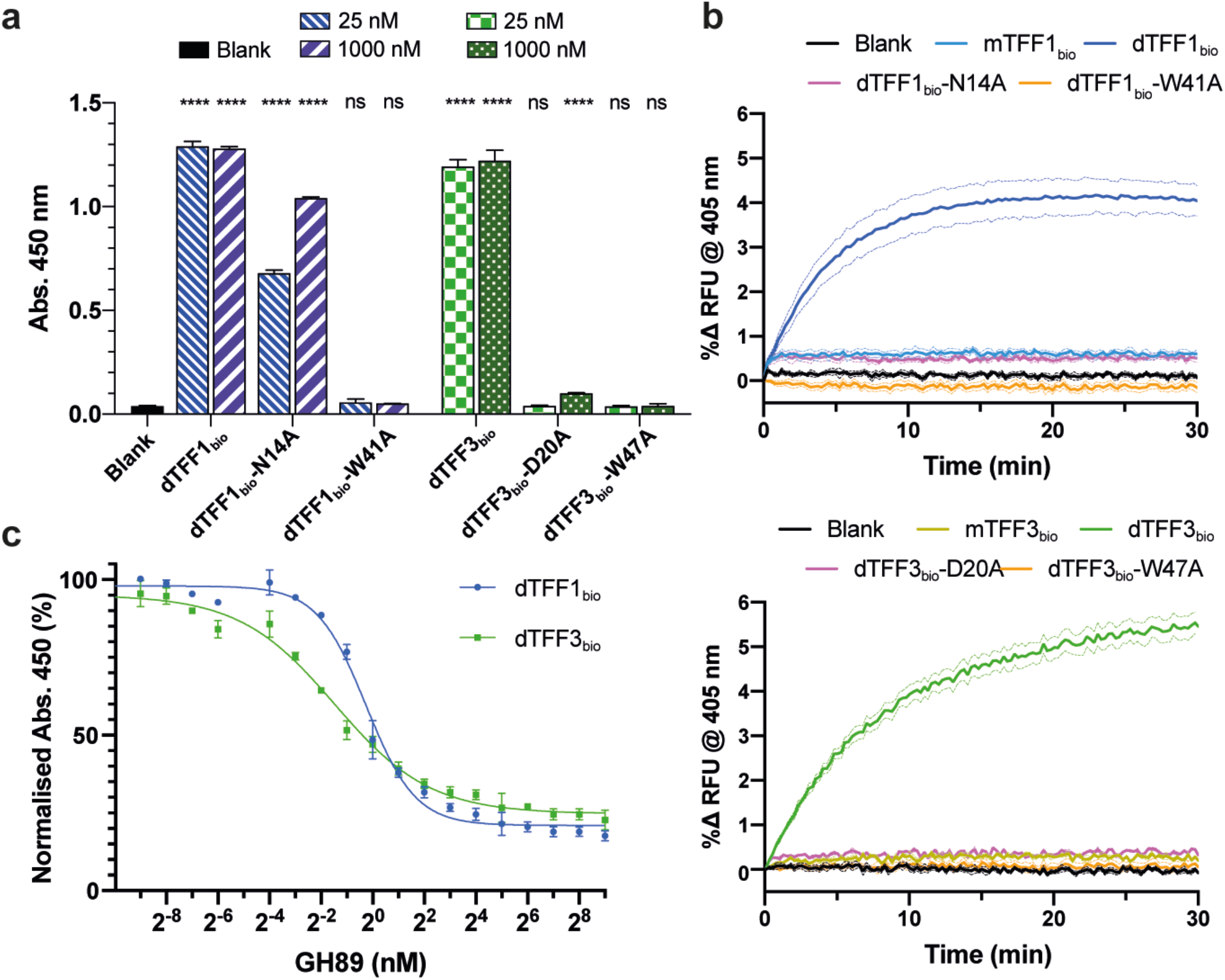
(a) The ability of dimeric WT or mutant dTFF1_bio_ (blue), or dTFF3_bio_ (green) to bind to immobilized pMucin_red_ assessed by ELISA. Error bars indicate standard deviations of three replicates. ns = p>0.05, **** = p<0.0001. (b) pMucin agglutination assays using monomeric wild-type, dimeric wild-type or mutant TFF1 (top) and TFF3 (bottom) with optical density at 405 nm monitored with respect to time. (c) *Cp*GH89-mediated displacement of dimeric TFF1 (blue circles) and TFF3 (green squares) from immobilized pMucin, as determined by ELISA. Error bars represent standard deviations from three replicates.

Mutagenesis of the ligand-binding Asn/Asp and Trp in dimeric TFF1 and TFF3 constructs (dTFF1_bio_ and dTFF3_bio_) enabled an examination of their role in mucin binding by ELISA. The dTFF1_bio_-N14A and dTFF3_bio_-D20A mutants had dramatically reduced affinity for pMucin_red_, while dTFF1_bio_-W41A and dTFF3_bio_-W47A had no detectable affinity for pMucin_red_ **(Fig. 2a)**. ITC was unable to detect any binding of GlcNAc-α-1,4-Gal to the TFF mutants (**Supplementary Fig. 1**), suggesting that the weak residual binding observed for dTFF1_bio_-N14A and dTFF3_bio_-D20A arises from avidity effects. Curiously, all known mammalian TFF1s utilise a disaccharide-binding Asn, while all TFF3s retain an Asp (**Supplementary Fig. 3**). TFF1 operates in the low pH gastric mucosa, while TFF3 does not, and so the Asn/Asp difference may relate to pH tolerances in glycan binding. To supplement binding data collected at pH 7.4 **(Fig. 1e)**, the Kd for the disaccharide–TFF complexes were determined at pH 5.0 and pH 2.6 using the same tryptophan fluorescence quenching assay **(Supplementary Fig. 4)**. At pH 2.6, the Kd for mTFF1 increased only slightly to 67 ± 1 μM, while for mTFF3 the Kd was 350 ± 60 μM; six-fold higher than at pH 7.4. Like TFF1, TFF2 is abundant in gastric mucus, has a conserved Asn (**Supplementary Fig. 3**) and also binds mucins at low pH^14^. The non-ionisable disaccharide-binding Asn side chain is thus important for TFF activity at low pH.

To demonstrate that the TFFs are capable of cross-linking mucosal glycoproteins into larger particles in a glycan-dependent manner, we performed agglutination assays using pMucin and our monomeric, dimeric and mutant TFF1/3 constructs **(Fig. 2b, Supplementary Fig. 5)**. Dimeric TFF1 and TFF3 induced a dose-dependent increase in light scattering over time at concentrations as low as 4 μM. Monomeric TFFs had no detectable influence on light scattering at concentrations up to 32 μM, commensurate with divalency being required for glycoprotein cross-linking. The mutant dimers, which had greatly diminished or no detectable disaccharide-binding activity, mirrored the results of the monomers in that they failed to agglutinate pMucins. We also demonstrated using ELISA that mucin-bound dTFF1_bio_ and dTFF3_bio_ could be displaced by the *Cp*GH89 enzyme-catalysed removal of α-GlcNAc **(Fig. 2c)**. The efficacy of *Cp*GH89 under these conditions differed between the two TFFs, with EC50 values of 0.8 ± 0.1 nM for TFF1 and 0.4 ± 0.1 nM for TFF3: the slightly greater resilience of TFF1 may arise from its higher affinity for the disaccharide. This enzyme may be a useful tool for perturbing TFF function in other systems.

Finally, we re-evaluated structures of other human proteins with trefoil domains in the PDB to uncover further evidence that these domains evolved as lectins in other contexts: the trefoil domain of human lysosomal a-glucosidase is bound to isomaltose (Glc-α-1,6-Glc) (PDB ID: 5KZW) (**Supplementary Fig. 6**). Phylogenetic analysis of all mammalian proteins with a trefoil domain revealed three clades of putative lectins associated with either amylose-processing enzymes, the GlcNAc-α-1,4-Gal-binding mucosal TFFs, or the glycoprotein-binding zona pellucida proteins **(Fig. 3)**. Our work provides a basis for probing the impact of these domains on the function of their respective proteins.

**Figure 3.**
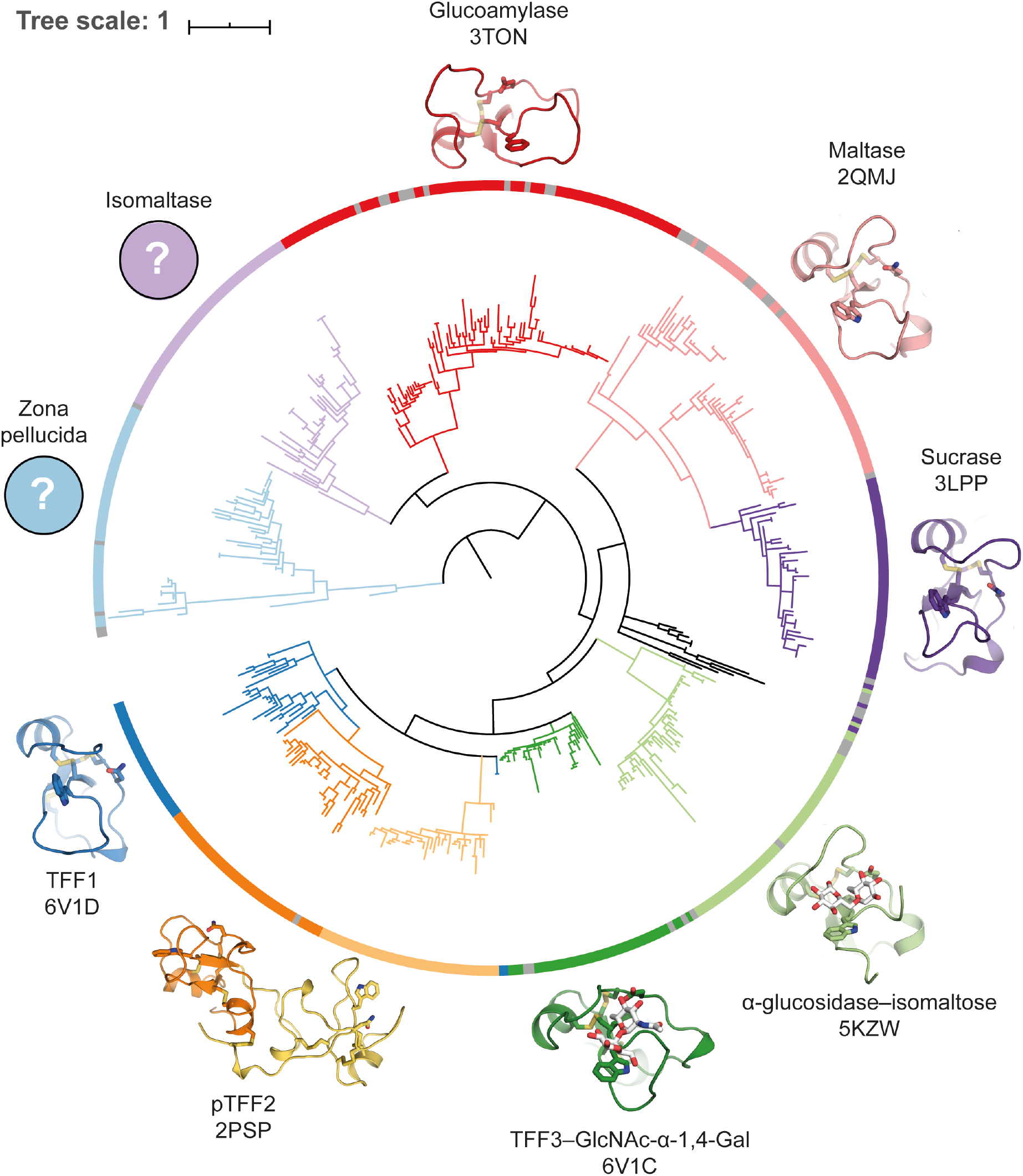
Phylogenetic tree of the mammalian trefoil domains color coded based on their gene- and enzyme-association, including structural representations where available (PDB Accession IDs listed under protein names). The large clade of trefoil domains found in amylose-processing enzymes likely bind maltose or isomaltose, as observed for lysosomal α-glucosidase. The clade of TFF trefoil domains bind the GlcNAc-α-1,4-Gal disaccharide that is unique to type-III mucin-like O-glycans. It is unclear what ligand the third clade, found in zona pellucida protein 1 and 4, might have affinity for.

The data presented here reveals that the mucin-binding activity of all TFFs is contingent on their glycoprotein binding-partners possessing α-GlcNAc-terminated O-glycans. These glycan structures are assembled in the Golgi by α-1,4-N-acetylglucosaminyltransferase (α4GnT)^16,17^, which is constitutively expressed in gastric mucous and Brunner’s gland cells^18^ but not in lung tissues. As such, future studies that focus on the role of the TFF lectins in health and disease must consider quantitation of α4GnT and α-GlcNAc-terminated O-glycans in cells and mucus, respectively, in addition to the abundance of the TFFs themselves.

## Acknowledgements

We thank A/Prof. Alisdair Boraston for providing the plasmid for *Cp*GH89 production^19^, Dr. Richard Birkinshaw for suggesting surface lysine methylation for protein crystallisation, the MX2 beamline staff at the Australian Synchrotron for help with X-ray data collection, as well as Dr. Janet Newman and Dr. Bevan Marshall at the Commonwealth Scientific and Industrial Research Organisation (CSIRO) Collaborative Crystallisation Centre (C3) for assistance in protein crystallization. This work was supported by a National Health and Medical Research Council of Australia (NHMRC) project grant (GNT1139549).

## Author contributions

M. J. performed all biophysical, bioinformatic and structural studies; M.J., J.P. and A.J. produced recombinant protein; N.E.S. performed all mass spectrometry experiments; E.D.G.-B. conceived the project; M.J. and E.D.G.-B. co-wrote the manuscript.

## Methods

### Cloning, expression and purification of untagged monomeric TFF1/3

A series of dsDNA oligonucleotides encoding human TFF1 (UniProt ID: P04155) and TFF3 (UniProt ID: Q07654) with an N-terminal PelB signal peptide, no interchain Cys, and a C-terminal His_6_-tag codon-harmonised for *E. coli* (**Supplementary Table 4)** were synthesised (IDT) and cloned into the pET29b(+) vector (Novagen) using the *NdeI* and *XhoI* restriction sites. The resulting plasmids were verified using Sanger sequencing. Each plasmid was transformed into T7 Express cells (NEB) and plated onto LB-agar + 2% glucose + 50 μg.ml^−1^ Kan and incubated at 37 °C for 16 h. Single colonies were picked to generate overnight cultures, which were used to inoculate SB media + 0.2% glucose + 50 μg.ml^−1^ Kan. The culture was incubated at 37°C and 220 rpm until it reached an OD600 of 1.0. IPTG was added to a final concentration of 0.1 mM for mTFF1 or 0.4 mM for mTFF3 and the culture was incubated for 4 h at 30°C and 220 rpm. The cells were harvested by centrifugation (8,000×g, 25 min, 4 °C) and frozen (−80°C) until further use.

Thawed cells were resuspended in ice-cold high-osmolyte buffer (0.5 M sucrose, 0.2 M Tris, pH 8.0) at 4°C for 15 min. Three volumes of ice-cold milliQ H_2_O was added and the mixture nutated at 4°C for 30 min. The suspension was adjusted to 150 mM NaCl, 2 mM MgCl_2_, 20 mM ImH, pH 8.0, centrifuged (18,000×g, 30 min, 4°C) and the supernatant filtered (0.45 μm). The supernatant was applied to a Ni-affinity column (HisTrap Excel, 1 ml), the column washed with 10 CV of 50 mM Tris, 300 mM NaCl, 40 mM ImH, pH 7.5, and the protein eluted with 50 mM Tris, 300 mM NaCl, 400 mM ImH, pH 7.5.

Fractions containing product, as judged by SDS-PAGE, were pooled and further purified by size exclusion chromatography (Superdex^®^ 75 10/300, GE Healthcare) using 50 mM Tris-HCl, 150 mM NaCl, pH 7.5 as buffer. The C-terminal His6-tag was removed by the addition of 1:100 molar ratio of carboxypeptidase A (Sigma) with incubation at 25 °C for 24 h. The solution was filtered (0.22 μM) and passed through a Ni-affinity column (HisTrap Excel, 1 ml) with the flow-through being further purified by size exclusion chromatography (Superdex^®^ 75 10/300, GE Healthcare) using 50 mM HEPES, 50 mM NaCl, pH 7.4 (ITC-Buffer). SDS-PAGE and intact ESIMS data confirmed the homogeneity of mTFF1 and mTFF3, the removal of the His_6_-tags and the presence of three disulfide bonds (**Supplementary Fig. 7 and 8**).

### Cloning, expression and purification of avi-tagged-TFF1/3 and their mutants

A series of dsDNA oligonucleotides encoding human TFF1 (UniProt ID: P04155), TFF3 (UniProt ID: Q07654) and mutants thereof with an N-terminal His_6_-Avi-tag and Factor Xa cleavage site codon-harmonised for *E. coli* (**Supplementary Table 4)** were synthesised (IDT) and cloned into the pET28b(+) vector (Novagen) using the *NcoI* and *XhoI* restriction sites. The resulting plasmids were verified using Sanger sequencing. Each plasmid was transformed into SHuffle T7 cells (NEB) and plated onto LB-agar + 2% glucose + 50 μg.ml^−1^ Kan and incubated at 37 °C for 16 h. Single colonies were picked to generate overnight cultures, which were used to inoculate SB media + 0.2% glucose + 50 μg.ml^−1^ Kan. The culture was incubated at 30°C and 220 rpm until it reached an OD_600_ of 1.0. IPTG was added to a final concentration of 0.4 mM and the culture was incubated for 4 h at 30°C and 220 rpm. The cells were harvested by centrifugation (8,000×g, 25 min, 4 °C) and frozen (−80°C) until further use.

The cell pellet from 1.0 l of culture was resuspended in 20 ml ice-cold PBS (50 mM NaP_i_, 150 mM NaCl, pH 7.5) + 40 mM ImH + 0.5 mM EDTA + 2 μL Benzonaze (Millipore) + 1 tablet of cOmplete EDTA-free protease inhibitor cocktail (Roche). The suspension was sonicated in an ice bath in bursts of 5 seconds, followed by 10 seconds pause, at 50% amplitude until total energy spent reached 10 kW. The lysate was centrifuged (18,000×g, 30 min, 4°C) and the supernatant filtered (0.45 μm). The supernatant was applied to a Ni-affinity column (HisTrap Excel, 1 ml), the column washed with 10 CV of 50 mM Tris, 300 mM NaCl, 40 mM ImH, pH 7.5, and the protein eluted with 50 mM Tris, 300 mM NaCl, 400 mM ImH, pH 7.5. Fractions containing product, as judged by SDS-PAGE, were pooled and further purified by size exclusion chromatography (Superdex^®^ 75 10/300, GE Healthcare) using 50 mM Tris-HCl, 150 mM NaCl, pH 7.5 as buffer. Non-reducing SDS-PAGE and intact ESI-MS data suggested that these proteins had one free cysteine each: the interchain disulfide had not formed within the cell. These were biotinylated (see below) to generate mTFF1_bio_ and mTFF3_bio_.

To make the native dimers with an interchain disulfide bond, protein samples at a concentration of 5 mg.ml^−1^ in 100 mM Tris-HCl, pH 8.3 were treated with 20 mM K_3_Fe(CN_6_) for 16 h at 25 °C. The reaction mixture was filtered (0.22 μm) and purified by size exclusion chromatography (Superdex^®^ 75 10/300, GE Healthcare) using 50 mM Tris-HCl, pH 7.5 as buffer, and anion exchange chromatography (MonoQ 5/50 GL, GE Healthcare) using a gradient over 30 CV from 100% binding buffer (50 mM Tris-HCl, pH 7.5) to 60% elution buffer (binding buffer + 1 M NaCl). SDS-PAGE and intact ESI-MS data confirmed the homogeneity and number of disulfide bonds in each sample (**Supplementary Fig. 7 and 8**).

### Cloning, expression and purification of TFF2

A dsDNA oligonucleotide encoding human TFF2 (UniProt ID: Q03403) with an N-terminal gp67 signal peptide and a C-terminal His10 sequence codon-harmonised for *Spodoptera frugiperda* (**Supplementary Table 4)** was synthesised (IDT) and cloned into the pFastBac vector (ThermoFisher) using the *SpeI* and *XhoI* restriction sites. The resulting plasmid was verified using Sanger sequencing. Expression of TFF2 was achieved in *Sf21* cells using the ‘Bac-to-Bac Baculovirus Expression System’ (ThermoFisher) in accordance with the manufacturer’s instructions. One litre of cell culture at a density of 1×10^6^ cells.ml^−1^ was infected with 30 ml of P3 baculovirus and cultured at 27 °C for 72 h. The culture was centrifuged (8,000×g, 20 min, 4 °C) and the supernatant collected, adjusted to pH 7.5 and filtered (0.45 μm). The supernatant was applied to a Ni-affinity column (HisTrap Excel, 5 ml), the column washed with 10 CV of 50 mM Tris, 300 mM NaCl, 40 mM ImH, pH 7.5, and the protein eluted with 50 mM Tris, 300 mM NaCl, 400 mM Imidazole, pH 7.5. Fractions containing product, as judged by SDS-PAGE, were pooled and further purified by size exclusion chromatography (Superdex^®^ 75 10/300, GE Healthcare) using 50 mM Tris-HCl, 150 mM NaCl, pH 7.5 as buffer. SDS-PAGE and intact ESI-MS data confirmed that TFF2 was homogenous and had seven disulfide bonds (**Supplementary Fig. 7 and 8**).

### Biotinylation of Avi-tagged TFFs

Biotinylation of Avi-tagged TFFs was accomplished by combining 952 μl of 100 μM TFF in PBS, 5 μl of 1 M MgCl_2_, 20 μl of 100 mM ATP, 20 μl of 40 μM BirA (Sigma) and 3 μl of 50 mM D-biotin. The solution was incubated for 1 h at 30°C with gentle agitation. An additional 20 μl of 100 mM ATP, 20 μl of 40 μM BirA and 3 μl of 50 mM D-biotin was added and the solution incubated for an additional hour. The sample was filtered (0.22 μm) and purified by size exclusion chromatography (Superdex^®^ 75 10/300, GE Healthcare) using 50 mM Tris-HCl, 150 mM NaCl, pH 7.5 as buffer. The degree of biotinylation was assessed by intact ESI-MS and was found to be near 100% in all cases (**Supplementary Fig. 8**).

### Preparation of mucin samples

A 50 mg.ml^−1^ stock of pMucin was prepared by resuspending porcine gastric mucin type III (Sigma) in PBS + 50 mM EDTA + 0.02% NaN_3_. This pMucin (1 ml) was dialyzed (100 kDa NMWL) against 4 M guanidium hydrochloride (2× 2.0 l) over 48 h at 4 °C. Precipitate was removed by centrifugation (8,000×g, 20 min, 4 °C) and the soluble supernatant dialyzed (100 kDa NMWL) against PBS (2× 2.0 l) over 48 h at 4 °C. Half of this sample was treated with 50 μg/mL *Cp*GH89 at 30°C for 16 h: the *Cp*GH89 was produced as previously described^19^. Both mucin samples were reduced with 10 mM DTT (30 min, 50°C) then alkylated by the addition of 30 mM iodoacetamide (30 min, 50°C). These samples were dialyzed (100 kDa NMWL) against PBS (2× 2.0 l) over 48 h at 4 °C. The concentration of the final pMucin_red_ and pMucin_red+GH89_ stock solutions were standardised to A_280_=1.5 using PBS. These samples were aliquoted, flash frozen in liquid N_2_ and stored at −80 °C.

### TFF-mucin ELISA

100 μl of either pMucin (10 μg.ml^−1^), 1:1000 pMucinred, or 1:1000 pMucin_red+GH89_ in 100 mM carbonate-bicarbonate buffer, pH 9.4, was added to each well of a 96-well plate (flat-bottom Nunc MaxiSorp, Thermo Scientific) and the plate gently nutated for 16 h at 4°C. The wells were rinsed four times with 200 μl PBS + 0.2% Tween (PBS-T) then blocked with 200 μL PBS + 5% (w/v) skim milk powder (blocking buffer) for 1 h at 25 °C. The wells were rinsed four times with 200 μl PBS-T then incubated with 50 μl blocking buffer + TFF (various concentrations) and 1:1000 Strep-HRP (Thermo) for 1 h at 25 °C. The wells were rinsed four times with 200 μl PBS-T then developed by adding 100 μl of TMB-Turbo (ThermoFisher) and incubated for 30 min at 25 °C. The reaction was quenched by the addition of 100 μl of 2 M H2SO4. Absorbance at 450 nm was read within 20 min using an EnVision 2105 Multimode Plate Reader (PerkinElmer). Data was plotted using Prism 8 (GraphPad).

### Isothermal calorimetry

All TFF and ligand stock solutions were prepared in the same buffer (50 mM HEPES, 50 mM NaCl, pH 7.4) and filtered (0.22 μm) prior to titration. All control and non-binding ligand experiments were performed as 12×3.18 μl injections while for GlcNAc-α-1,4-Gal, 25×1.58 μL injections were used. All runs were performed at 25 °C in a MicroCal iTC200 (Malvern).

### Tryptophan fluorescence quenching assay

Tryptophan fluorescence quenching assays were performed at 25 °C on an EnVision 2105 Multimode Plate Reader (PerkinElmer) with excitation at 280 nm and emission measured at 340 nm. The assay was performed in a 384-well plate (Corning, 384-well low volume black round bottom polystyrene non-binding surface) using 15 μl per well. Each sample contained mTFF1 or mTFF3 (5 μM) and GlcNAc-α-1,4-Gal (1 μM–4.1 mM) in 50 mM BTP/citric acid, 25 mM NaCl, at either pH 2.6, 5.0, or 7.4. For each dilution series, K_d_ was calculated by fitting the data to a one-site binding curve in Prism 8 (GraphPad) and the mean calculated from three independent experiments.

### Crystallisation and structure determination of apo-mTFF1

Sitting drops comprised of 1 μl well solution (1 M ammonium sulfate, 0.1 M Tris-HCl, pH 8.5) and 1 μL mTFF1 solution (10 mg.ml^−1^) supplemented with GlcNAc-α-1,4-Gal (5 mM) afforded small clusters of rod-like crystals after one month at 20°C. The crystals were cryo-protected by supplementing the mother liquor with 3 M ammonium sulfate before being collected on a crystal loop and cryogenically stored in liquid nitrogen. Data was collected at the Australian Synchrotron (MX2 beamline)^20^ and processed using XDS^21^. The structure was solved by molecular replacement using PHASER^22^ and a truncated version of the mTFF3 crystal structure described below as a search model. No density commensurate with a GlcNAc-α-1,4-Gal ligand was observed. The final model of TFF1 in its apo-form was built in Coot^23^ and refined with Phenix^24^ to a resolution of 2.4 Å. Data collection and refinement statistics are summarized in **Supplementary Table 3**. The Rwork and R_free_ after the final refinement was 0.1939 and 0.2259, respectively. The coordinates have been deposited in the Protein Data Bank (accession code: 6V1D). Figures were prepared using PyMOL.

### Crystallisation and structure determination of mTFF3 in complex with GlcNAc-α-1,4-Gal

Surface lysine methylation of mTFF3 was conducted using the protocol described by Kim *et al*^25^. Sitting drops comprised of 0.15 μL well solution (2 M ammonium sulfate, 0.2 M potassium sodium tartrate, and 0.1 M trisodium citrate-citric acid, pH 5.6) and 0.15 μL of permethylated mTFF3 solution (15 mg.ml^−1^) supplemented with GlcNAc-α-1,4-Gal (5 mM) afforded a single crystal after 14 days at 20°C. The crystal was cryo-protected by supplementing the mother liquor with 3 M sodium malonate-malonic acid, pH 5.6, before being collected on a crystal loop and cryogenically stored in liquid nitrogen. Data was collected at the Australian Synchrotron (MX2 beamline)^20^ and processed using XDS^21^. The structure was solved by molecular replacement using PHASER^22^ with a truncated version of the porcine TFF2 crystal structure as a search model (PDB ID: 2PSP)^26^. The final model was built in Coot^23^ and refined with Phenix^24^ to a resolution of 1.55 Å. Data collection and refinement statistics are summarized in **Supplementary Table 3**. The R_work_ and R_free_ after the final refinement was 0.1738 and 0.1882, respectively. The coordinates have been deposited in the Protein Data Bank (accession code: 6V1C). Figures were prepared using PyMol and LigPlot+^27^.

### Intact protein MS analysis

Intact analysis was performed using a 6520 Accurate mass Q-TOF mass spectrometer (Agilent). Protein samples were resuspended in 2% acetonitrile, 0.1% TFA and 5 μg loaded onto a C5 Jupiter column (5 μm, 300 Å, 50 mm × 2.1 mm, Phenomenex) using an Agilent 1200 series HPLC system. Samples were desalted by washing the column with buffer A (2% acetonitrile, 0.1% formic acid) for 4 min and then separated with a 12 min linear gradient from 2 to 100% buffer B (80% acetonitrile, 0.1% formic acid) at a flow rate of 0.250 ml.min^−1^. MS1 mass spectra were acquired at 1 Hz between a mass range of 300–3,000 *m/z*. Intact mass analysis and deconvolution was performed using MassHunter B.06.00 (Agilent).

### Mucin agglutination assay

Mucin agglutination assays were performed at 25 °C on an EnVision 2105 Multimode Plate Reader (PerkinElmer) to monitor transmission at 405 nm over time. The assay was performed in a 96-well plate (Corning, 96-well round bottom polystyrene non-binding surface) with 25 μl per well. pMucin concentration was kept consistent at 0.02% w/v and TFF concentration varied (0, 1, 2, 4, 8, 16, or 32 μM). The resulting curves were normalized to percent decrease in signal, averaged over three replicates and plotted using Prism 8 (GraphPad).

### TFF displacement assay

100 μl of pMucin (5 μg.ml^−1^) in 100 mM carbonate-bicarbonate buffer, pH 9.4, was added to each well of a 96-well plate (flatbottom Nunc MaxiSorp, Thermo Scientific) and the plate gently nutated for 1 h at 30°C. The wells were rinsed four times with 200 μl PBS + 0.2% Tween (PBS-T) then blocked with 200 μL PBS + 5% (w/v) skim milk powder (blocking buffer) for 1 h at 30 °C. The wells were rinsed four times with 200 μl PBS-T then incubated with 50 μl blocking buffer + dTFF1/3_bio_ (16 nM) for 1 h at 25 °C. The wells were rinsed four times with 200 μl PBS-T then treated with 100 μl GH89 (0-512 nM) in blocking buffer for 20 h at 30 °C. The wells were rinsed six times with 200 μl PBS-T then probed with 100 μl Strep-HRP diluted 1:1000 in blocking buffer for 1 h at 25 °C. The wells were rinsed six times with 200 μl PBS-T then developed by adding 100 μl of TMB-Turbo (ThermoFisher) and incubated for 30 min at 25 °C. The reaction was quenched by the addition of 100 μl of 2 M H_2_SO_4_. Absorbance at 450 nm was read within 20 min using an EnVision 2105 Multimode Plate Reader (PerkinElmer) and an EC_50_ calculated using Prism 8 (GraphPad) and the mean from three independent experiments

### Trefoil domain sequence analysis

An annotated sequence alignment of the trefoil domains of human TFF1, TFF2, and TFF3, was created by ESPript (espript.ibcp.fr)^28^. Sequence logos for each of the four mucosal TFF domains in the three human trefoil factors were generated by WebLogo (weblogo.berkeley.edu)^29^ using full length protein sequences annotated as mammalian TFF1, TFF2, or TFF3 retrieved from the UniProt database^30^ and aligned with Clustal Omega^31^. The phylogenetic tree of all mammalian trefoil domains was created as follows. All trefoil domain sequences corresponding to PF00088 were retrieved aligned according to their hidden Markov model logo generated by the Pfam database^32^. Based on UniProt^30^ metadata entries marked as obsolete or truncated were discarded. The remaining sequences were annotated as belonging to one of ten groups (glucoamylase, isomaltase, lysosomal α-glucosidase, maltase, sucrase, TFF1, TFF2-1, TFF2-2, TFF3, or zona pellucida) based on their UniProt annotation. The remaining alignment was used as input to calculate phylogenetic distances with phylogeny.fr^33^. The phylogenetic tree was visualized with iTol^34^ and colour-coded according to the described metadata. Where available, a structural representation of each clade of TFF-domains was created using PyMOL and included for eight of the ten groups mentioned above.

