## Supplementary Information for "Trefoil factors share a lectin activity that defines their role in mucus"

### SUPPLEMENTARY RESULTS

#### Supplementary Tables

**Supplementary Table 1.** Protein constructs used in this study (mutations are in red, biotinylated Lys in green and Cys forming interchain disulfides in yellow).

| construct | protein sequence |
|---|---|
| mTFF1 | QTETCTVAPRERQNCGFPGVTPSQCANKGCCFDDTVRGVPWCFYPNTILEK |
| mTFF1$_{\mathrm{bio}}$ | MDHHHHHHGLNDIFEAQKIEWHEIDGRQTETCTVAPRERQNCGFPGVTPSQCANKGCCF<br>DDTVRGVPWCFYPNTIDVPPEEECEF |
| dTFF1$_{\mathrm{bio}}$ | MDHHHHHHGLNDIFEAQKIEWHEIDGRQTETCTVAPRERQNCGFPGVTPSQCANKGCCF<br>DDTVRGVPWCFYPNTIDVPPEEECEF |
| dTFF1$_{\mathrm{bio}}$-N14A | MDHHHHHHGLNDIFEAQKIEWHEIDGRQTETCTVAPRERQACGFPGVTPSQCANKGCCF<br>DDTVRGVPWCFYPNTIDVPPEEECEF |
| dTFF1$_{\mathrm{bio}}$-W41A | MDHHHHHHGLNDIFEAQKIEWHEIDGRQTETCTVAPRERQNCGFPGVTPSQCANKGCCF<br>DDTVRGVPACFYPNTIDVPPEEECEF |
| TFF2 | EKPSPCQCSRLSPHQRTNCGFPGITSDQCFDNGCCFDSSVTGVPWCFHPLPKQESDQCV<br>MEVSDRRNCGYPGISPEECASRKCCFSNFIFEVPWCFFPKSVEDCHYKHHHHHHHHHH |
| mTFF3 | HHHHHHGLNDIFEAQKIEWHEIDGRGSEEYVGLSANQCAVPAKDRVDCGYPHVTPKECN<br>NRGCCFDSRIPGVPWCFKPLQEAEST |
| mTFF3$_{\mathrm{bio}}$ | MDHHHHHHGLNDIFEAQKIEWHEIDGREEYVGLSANQCAVPAKDRVDCGYPHVTPKECN<br>NRGCCFDSRIPGVPWCFKPLQEAECTF |
| dTFF3$_{\mathrm{bio}}$ | MDHHHHHHGLNDIFEAQKIEWHEIDGREEYVGLSANQCAVPAKDRVDCGYPHVTPKECN<br>NRGCCFDSRIPGVPWCFKPLQEAECTF |
| dTFF3$_{\mathrm{bio}}$-D20A | MDHHHHHHGLNDIFEAQKIEWHEIDGREEYVGLSANQCAVPAKDRVACGYPHVTPKECN<br>NRGCCFDSRIPGVPWCFKPLQEAECTF |
| dTFF3$_{\mathrm{bio}}$-W47A | MDHHHHHHGLNDIFEAQKIEWHEIDGREEYVGLSANQCAVPAKDRVDCGYPHVTPKECN<br>NRGCCFDSRIPGVPACFKPLQEAECTF |

**Supplementary Table 2.** Parameters used for ITC and thermodynamic terms determined for GlcNAc-α-1,4-Gal binding to the TFFs. Errors are listed as standard deviations (n=3).

| **Protein (30 μM)** | **GlcNAc-α-1,4-Gal (μM)** | **T (°C)** | **n** | **$K_d$ (μM)** | **ΔH (kcal · $mol^{-1}$)** | **ΔS (cal · $mol^{-1}$ · $K^{-1}$)** | **-TΔS (kcal · $mol^{-1}$)** | **ΔG (kcal · $mol^{-1}$)** |
|---|---|---|---|---|---|---|---|---|
| mTFF1 | 1250 | 25 | 1.0 ± 0.2 | 49 ± 4 | -10 ± 2 | -14 ± 5 | 4 ± 2 | -5.97 ± 0.03 |
| TFF2 | 1250 | 25 | 0.7 ± 0.3 | 44 ± 8 | -20 ± 10 | -60 ± 40 | 20 ± 10 | -6.2 ± 0.2 |
| mTFF3 | 1250 | 25 | 1.0 ± 0.1 | 65 ± 5 | -10.8 ± 0.5 | -17 ± 2 | 5.0 ± 0.5 | -5.83 ± 0.05 |

**Supplementary Table 3.** Refinement statistics for the structures reported in this study.

| Structure | HsTFF1 (PDB:6V1D) | HsTFF3:α-GlcNAc (PDB:6V1C) |
|---|---|---|
| Space group | P 1 21 1 | P 41 21 2 |
| No of protein chains in AU | 3 | 1 |
| Cell dimensions | | |
| *a*, *b*, *c* (Å) | 44.93, 41.90, 45.83 | 37.59, 37.59, 87.48 |
| a, b, g (°) | 90, 115.57, 90 | 90, 90, 90 |
| Wavelength (Å) | 0.9537 | 0.9537 |
| Resolution (Å)* | 41.34-2.40 (2.49-2.40) | 43.74-1.55 (1.58-1.55) |
| $R_{sym}$ or $R_{merge}$* | 0.231 (0.782) | 0.081 (1.949) |
| $R_{pim}$* | 0.225 (0.759) | 0.035 (0.849) |
| *I* / s*I** | 2.7 (1.0) | 15.8 (1.3) |
| CC(1/2) | 0.958 (0.570) | 0.999 (0.526) |
| Completeness (%)* | 98.6 (97.3) | 100 (100) |
| Redundancy* | 3.3 (3.4) | 11.5 (11.5) |
| Wilson B-factor (Å$^2$) | 8.2 | 20.3 |
| **Refinement** | | |
| Resolution (Å) | 41.34-2.40 | 34.54-1.55 |
| No. reflections | 6,046 | 9,706 |
| $R_{work}$ / $R_{free}$ | 0.1939 / 0.2259 | 0.1738 / 0.1882 |
| No. non-hydrogen atoms | | |
| Protein | 1114 | 404 |
| GlcNAc-α-1,4-Gal | n.a | 26 |
| Water | 96 | 38 |
| *B*-factors | | |
| Protein | 19.7 | 27.0 |
| GlcNAc-α-1,4-Gal | n.a | 26.4 |
| Water | 21.5 | 37.6 |
| R.m.s. deviations | | |
| Bond lengths (Å) | 0.002 | 0.009 |
| Bond angle (°) | 0.604 | 1.155 |
| Ramachandran plot (%) | | |
| Favored | 97.81 | 97.62 |
| Allowed | 2.19 | 2.38 |
| Disallowed | 0 | 0 |

* Values in paranthesis indicate values for highest resolution shell

**Supplementary Table 4.** Synthetic dsDNA used to prepare expression plasmids in this study (restriction sites underlined).

| sequence name | nucleotide sequence |
|---|---|
| mTFF1 | AAAAACATATGAAATACCTGCTGCCGACCGCTGCTGCTGGTCTGCTGCTCCTCGCTGCCCAGCCGGCGATGGCC<br>CAGACCGAAACATGTACCGTAGCACCTAGAGAACGCCAGAACTGCGGCTTCCCGGGCGTGACGCCGTCACAATG<br>TGCTAATAAAGGCTGCTGTTTTGATGATACCGTTAGAGGCGTACCGTGGTGCTTCTATCCGAACACCATTCTCG<br>AGAAAAA |
| aviTFF1 | AAAAACCATGGATCACCACCACCACCACCACGGTCTGAACGACATCTTCGAAGCGCAGAAGATCGAATGGCACG<br>AGATTGATGGCCGTCAAACCGAAACCTGCACCGTGGCGCCGCGTGAGCGTCAGAACTGCGGTTTTCCGGGCGTT<br>ACCCCGAGCCAATGCGCGAACAAAGGTTGCTGCTTCGACGATACCGTGCGTGGCGTTCCGTGGTGCTTTTACCC<br>GAACACCATTGACGTGCCGCCGGAGGAAGAGTGCGAGTTCTAACTCGAGAAAAA |
| aviTFF1-N14A | AAAAACCATGGATCACCACCACCACCACCACGGTCTGAACGACATCTTCGAAGCGCAGAAGATCGAATGGCACG<br>AGATTGATGGCCGTCAAACCGAAACCTGCACCGTGGCGCCGCGTGAGCGTCAGGCCTGCGGTTTTCCGGGCGTT<br>ACCCCGAGCCAATGCGCGAACAAAGGTTGCTGCTTCGACGATACCGTGCGTGGCGTTCCGTGGTGCTTTTACCC<br>GAACACCATTGACGTGCCGCCGGAGGAAGAGTGCGAGTTCTAACTCGAGAAAAA |
| dTFF1$_{bio}$-W41A | AAAAACCATGGATCACCACCACCACCACCACGGTCTGAACGACATCTTCGAAGCGCAGAAGATCGAATGGCACG<br>AGATTGATGGCCGTCAAACCGAAACCTGCACCGTGGCGCCGCGTGAGCGTCAGAACTGCGGTTTTCCGGGCGTT<br>ACCCCGAGCCAATGCGCGAACAAAGGTTGCTGCTTCGACGATACCGTGCGTGGCGTTCCGGCGTGCTTTTACCC<br>GAACACCATTGACGTGCCGCCGGAGGAAGAGTGCGAGTTCTAACTCGAGAAAAA |
| TFF2 | AAAAAACTAGTAAATCAGTCACACCAAGGCTTCAATAAGGAACACACAAGCAAGATGGTAAGCGCTATTGTTTT<br>ATATGTGCTTTTGGCGGCGGCGGCGCATTCTGCCTTTGCGGAAAAGCCTTCTCCTTGTCAATGCTCTCGTCTGT<br>CTCCTCACCAACGTACCAACTGCGGTTTCCCTGGTATCACCTCTGATCAATGCTTCGACAACGGTTGCTGCTTC<br>GACTCTTCTGTGACCGGTGTGCCATGGTGCTTCCACCCTCTGCCTAAGCAAGAATCTGACCAATGCGTGATGGA<br>AGTGTCTGACCGTCGTAACTGCGGTTACCCTGGTATCTCTCCTGAAGAATGCGCTTCTCGTAAGTGCTGCTTCT<br>CTAACTTCATCTTCGAAGTGCCTTGGTGCTTCTTCCCTAAGTCTGTGGAAGACTGCCACTACAAACACCATCAT<br>CACCACCATCACCACCATCACTGACTCGAGAAAAA |
| mTFF3 | AAAAACATATGAAATACCTGCTGCCGACCGCTGCTGCTGGTCTGCTGCTCCTCGCTGCCCAGCCGGCGATGGCC<br>GAAGAATATGTAGGACTATCAGCTAACCAGTGTGCGGTACCGGCGAAAGATCGCGTTGATTGCGGTTATCCGCA<br>TGTGACGCCGAAAGAATGTAACAACCGCGGCTGCTGCTTTGATAGCCGTATTCCGGGCGTGCCGTGGTGTTTTA<br>AACCGCTGCAGGAAGCGGAAAGCACCCTCGAGAAAAA |
| aviTFF3 | AAAAACCATGGATCACCACCACCACCACCACGGTCTGAACGACATCTTCGAGGCGCAGAAGATCGAGTGGCACG<br>AAATTGATGGTCGTGAGGAATACGTTGGTCTGAGCGCGAACCAATGCGCGGTGCCGGCGAAGGACCGTGTTGAT<br>TGCGGCTATCCGCACGTGACCCCGAAAGAATGCAACAACCGTGGTTGCTGCTTTGACAGCCGTATTCCGGGCGT<br>TCCGTGGTGCTTCAAACCGCTGCAGGAAGCGGAATGCACCTTTTAACTCGAGAAAAA |
| aviTFF3-D20A | AAAAACCATGGATCACCACCACCACCACCACGGTCTGAACGACATCTTCGAGGCGCAGAAGATCGAGTGGCACG<br>AAATTGATGGTCGTGAGGAATACGTTGGTCTGAGCGCGAACCAATGCGCGGTGCCGGCGAAGGACCGTGTTGCT<br>TGCGGCTATCCGCACGTGACCCCGAAAGAATGCAACAACCGTGGTTGCTGCTTTGACAGCCGTATTCCGGGCGT<br>TCCGTGGTGCTTCAAACCGCTGCAGGAAGCGGAATGCACCTTTTAACTCGAGAAAAA |
| aviTFF3-W47A | AAAAACCATGGATCACCACCACCACCACCACGGTCTGAACGACATCTTCGAGGCGCAGAAGATCGAGTGGCACG<br>AAATTGATGGTCGTGAGGAATACGTTGGTCTGAGCGCGAACCAATGCGCGGTGCCGGCGAAGGACCGTGTTGAT<br>TGCGGCTATCCGCACGTGACCCCGAAAGAATGCAACAACCGTGGTTGCTGCTTTGACAGCCGTATTCCGGGCGT<br>TCCGGCGTGCTTCAAACCGCTGCAGGAAGCGGAATGCACCTTTTAACTCGAGAAAAA |

#### Supplementary Figures

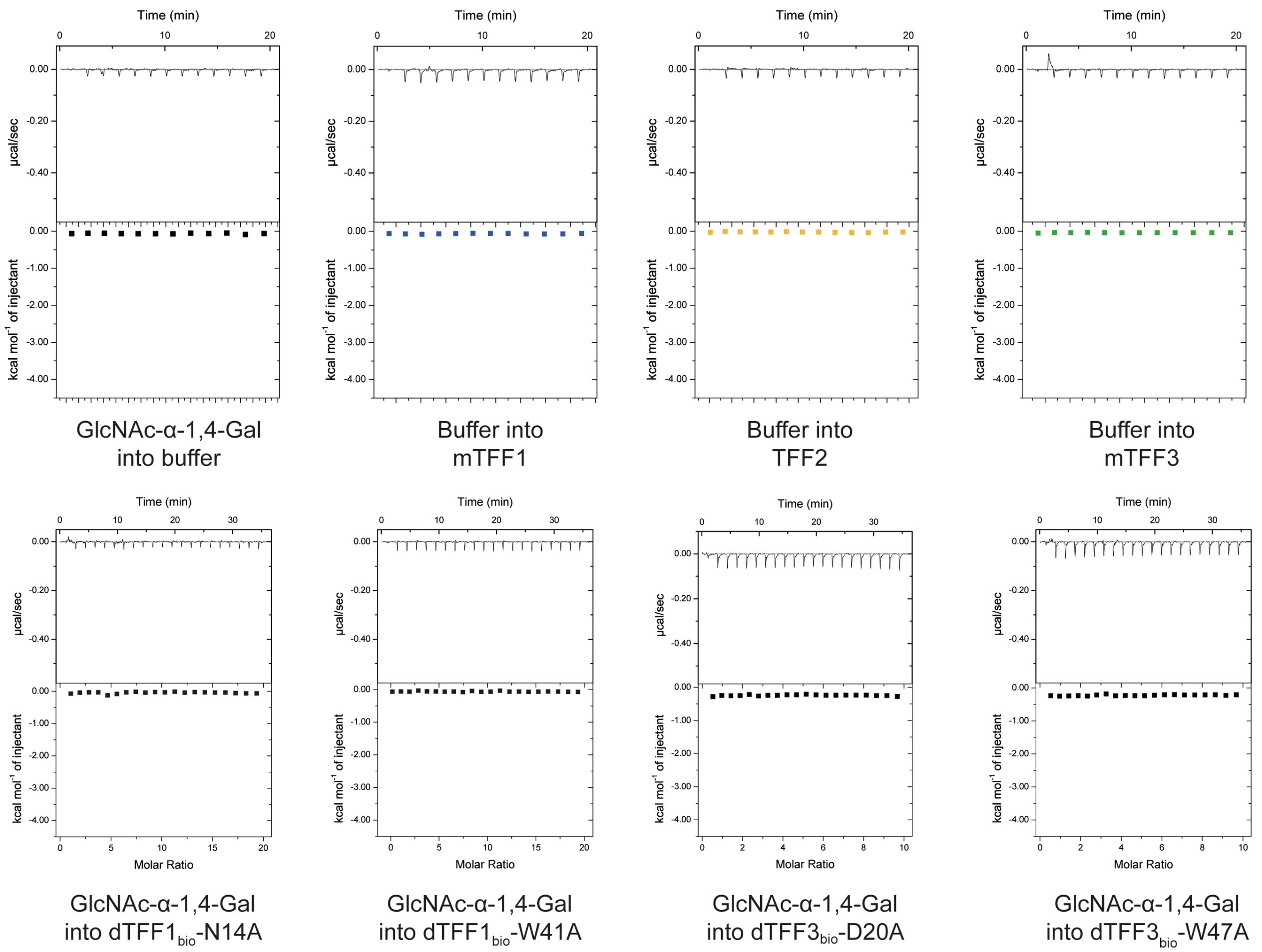

**Supplementary Figure 1.** Relevant controls for ITC.

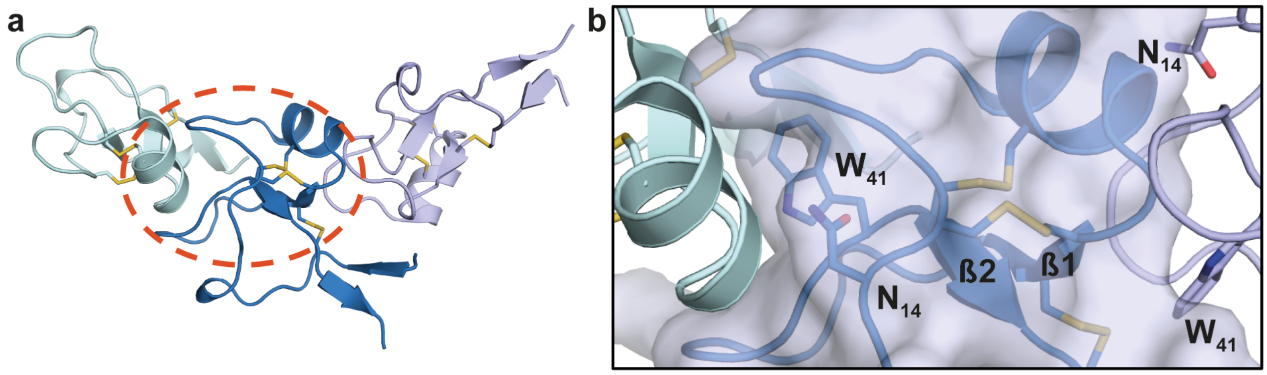

**Supplementary Figure 2.** The apo mTFF1 crystal structure has **a)** three molecules in the asymmetric unit and **b)** crystallographic contacts that occupy the ligand binding site.

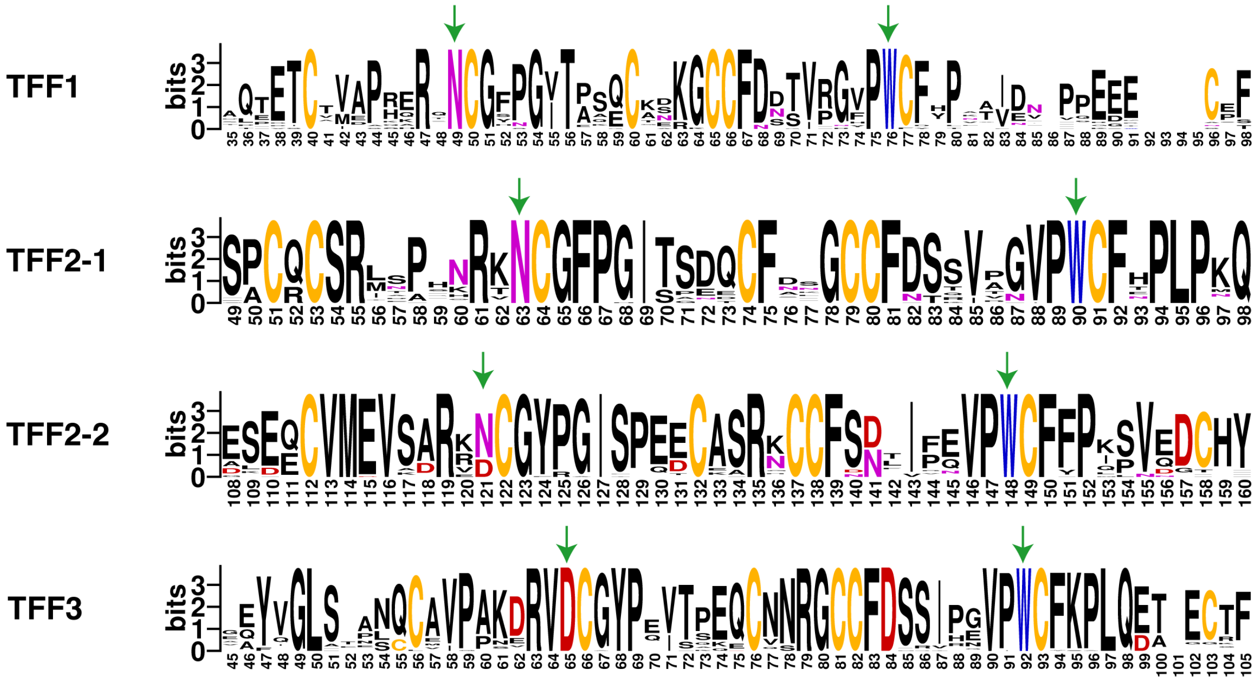

**Supplementary Figure 3.** Web-logos illustrating sequence conservation in the trefoil domains of mammalian TFF1-3. The residues with side-chains that interact with the GlcNAc-α-1,4-Gal ligand are indicated with green arrows.

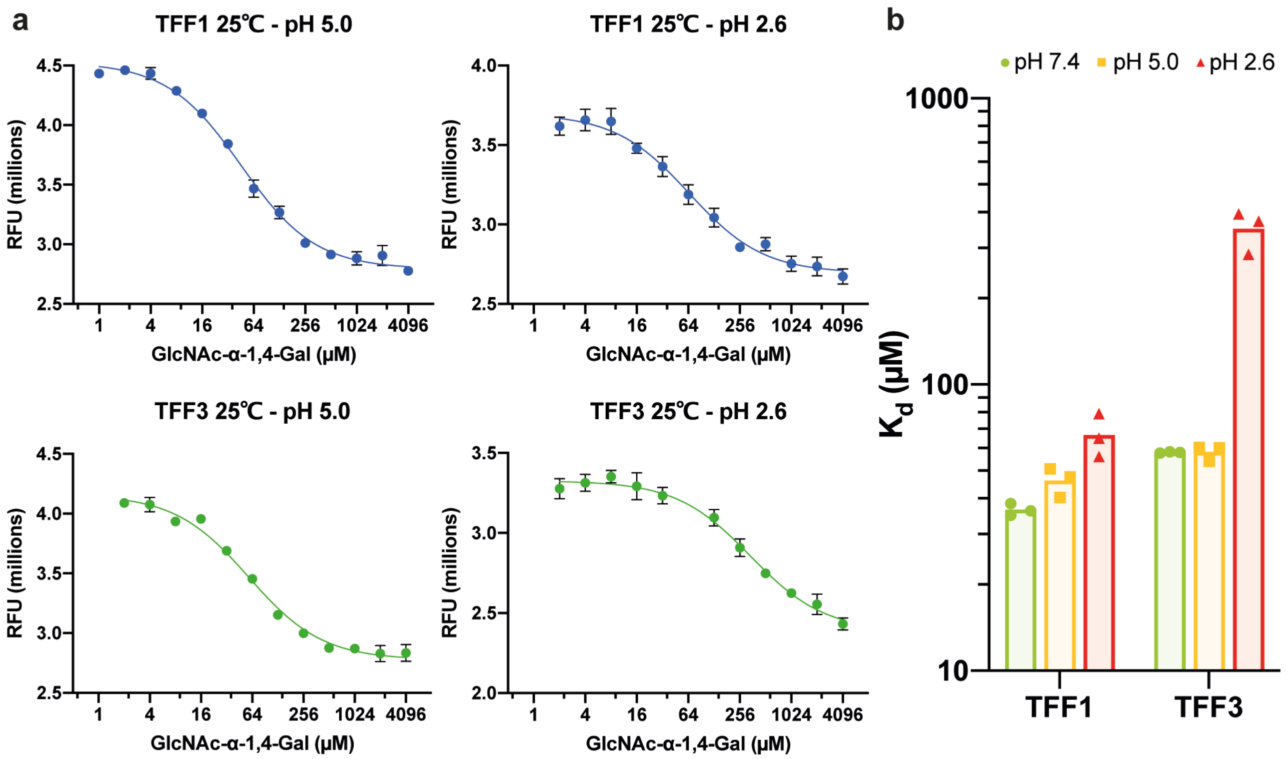

**Supplementary Figure 4.** Tryptophan fluorescence quenching assays at different pH for GlcNAc-α-1,4-Gal binding to mTFF1 and mTFF3. **a)** The titration curves for pH 5.0 and for pH 2.6 with the one-site curve fit. Errors indicate standard deviations of three replicates. **b)** Bar-graph of the average and individual log($K_d$) for each pH and TFF.

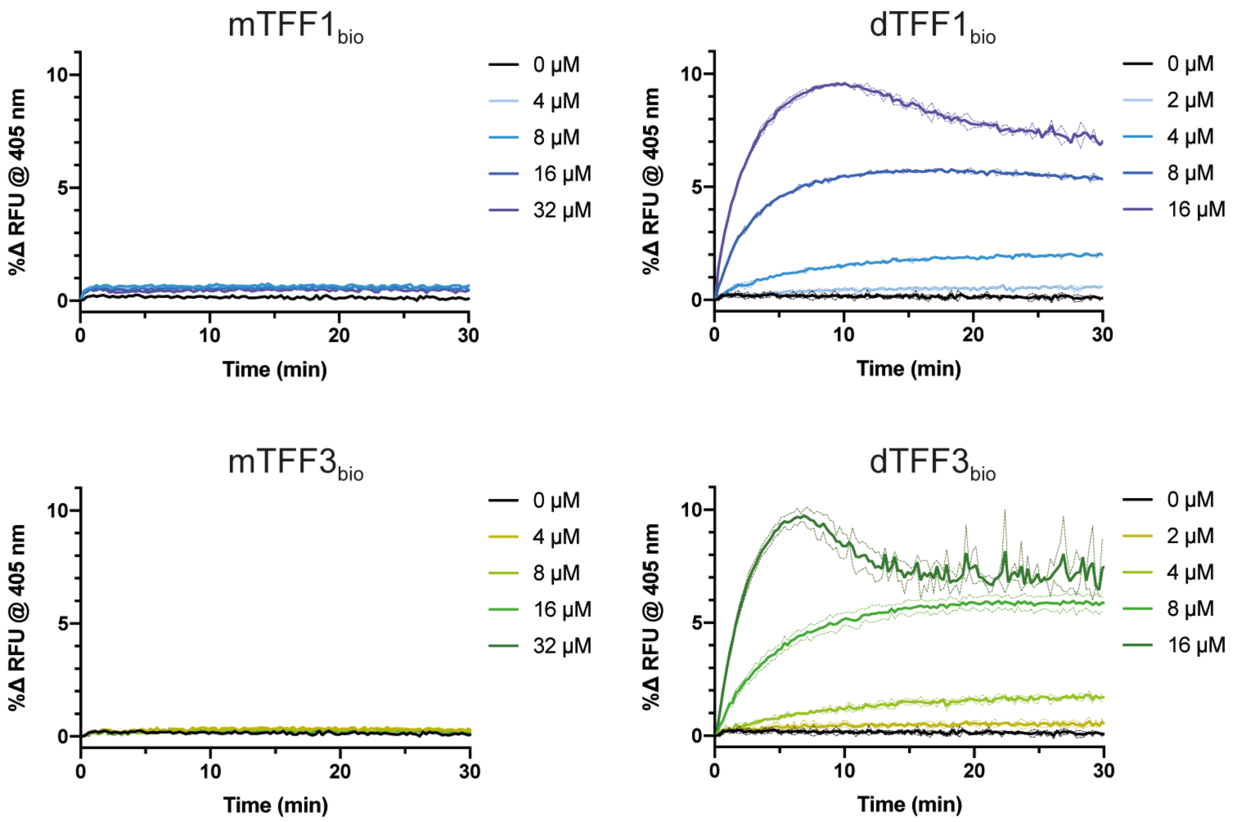

**Supplementary Figure 5.** Agglutination of pMucin at different concentrations of monomeric and dimeric TFF1 and TFF3. Curves are averaged from three replicates and dotted lines indicate standard deviations.

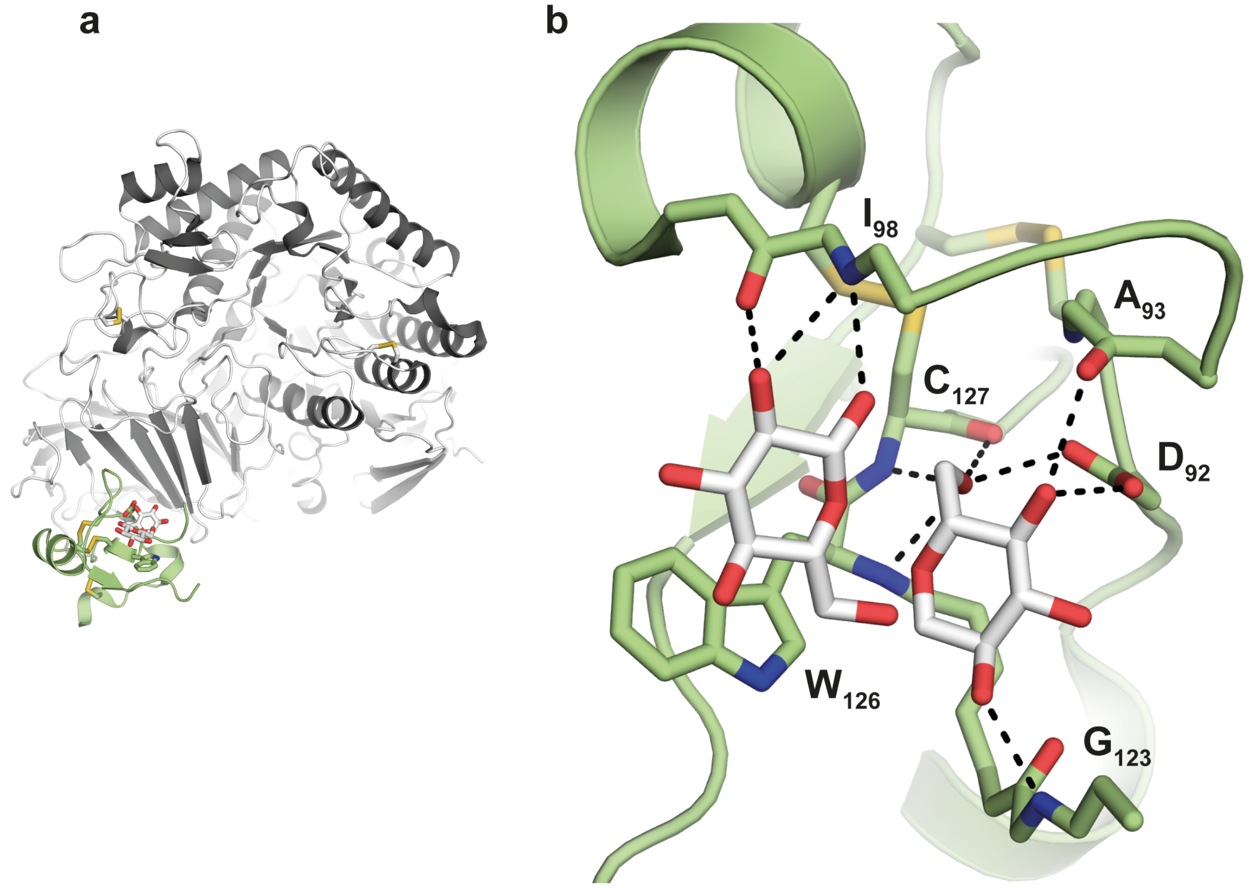

**Supplementary Figure 6. a)** The N-terminal trefoil domain (green cartoon) in human lysosomal α-glucosidase (dark grey cartoon) (PDB ID: 5KZW) is bound to isomaltose (white sticks with oxygen atoms coloured red). b) Close-up of the trefoil isomaltose binding site with residues forming hydrogen bonds drawn in sticks (sulfur, nitrogen and oxygen coloured yellow, blue and red, respectively).

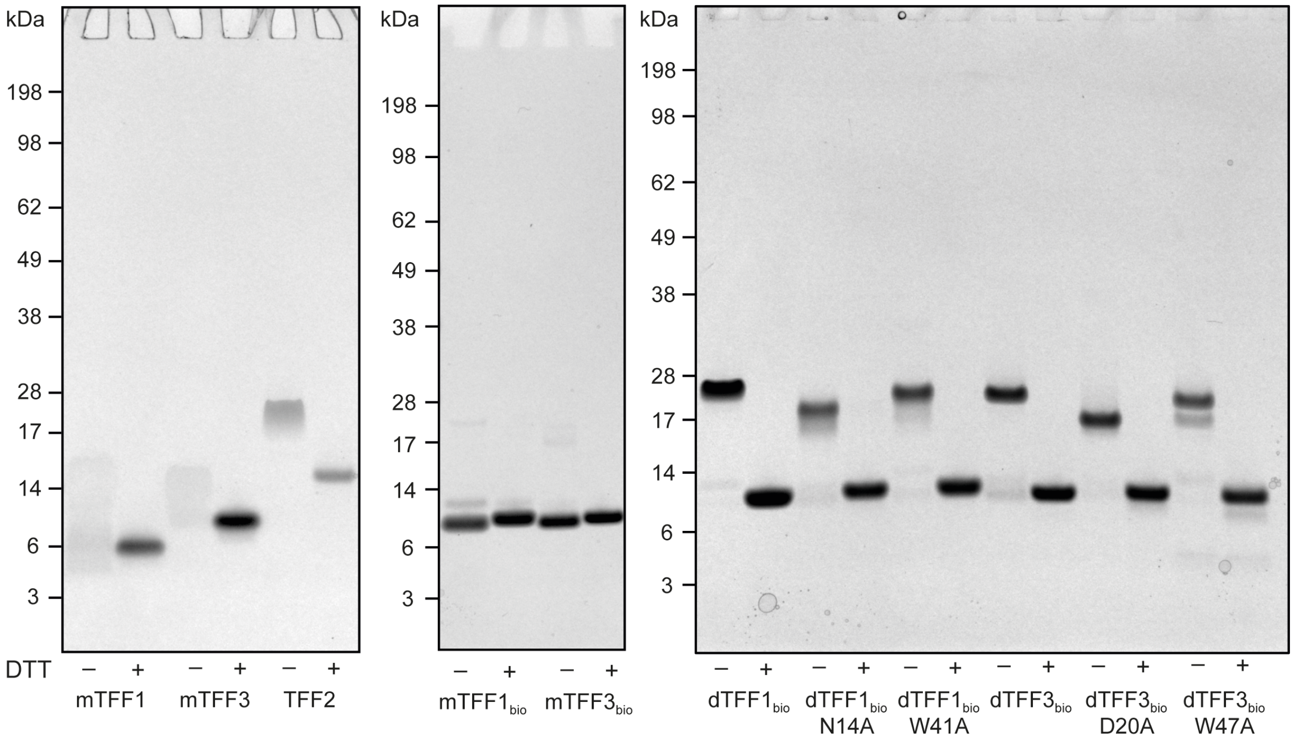

**Supplementary Figure 7.** Non-reducing and reducing SDS-PAGE of monomeric TFF1/3 and TFF2 (mTFF1, mTFF3, TFF2); biotinylated monomeric TFF1/3 ($mTFF1_{bio}$, $mTFF3_{bio}$); and biotinylated dimeric TFF1/3 ($dTFF1_{bio}$ and $dTFF3_{bio}$) and mutants thereof.

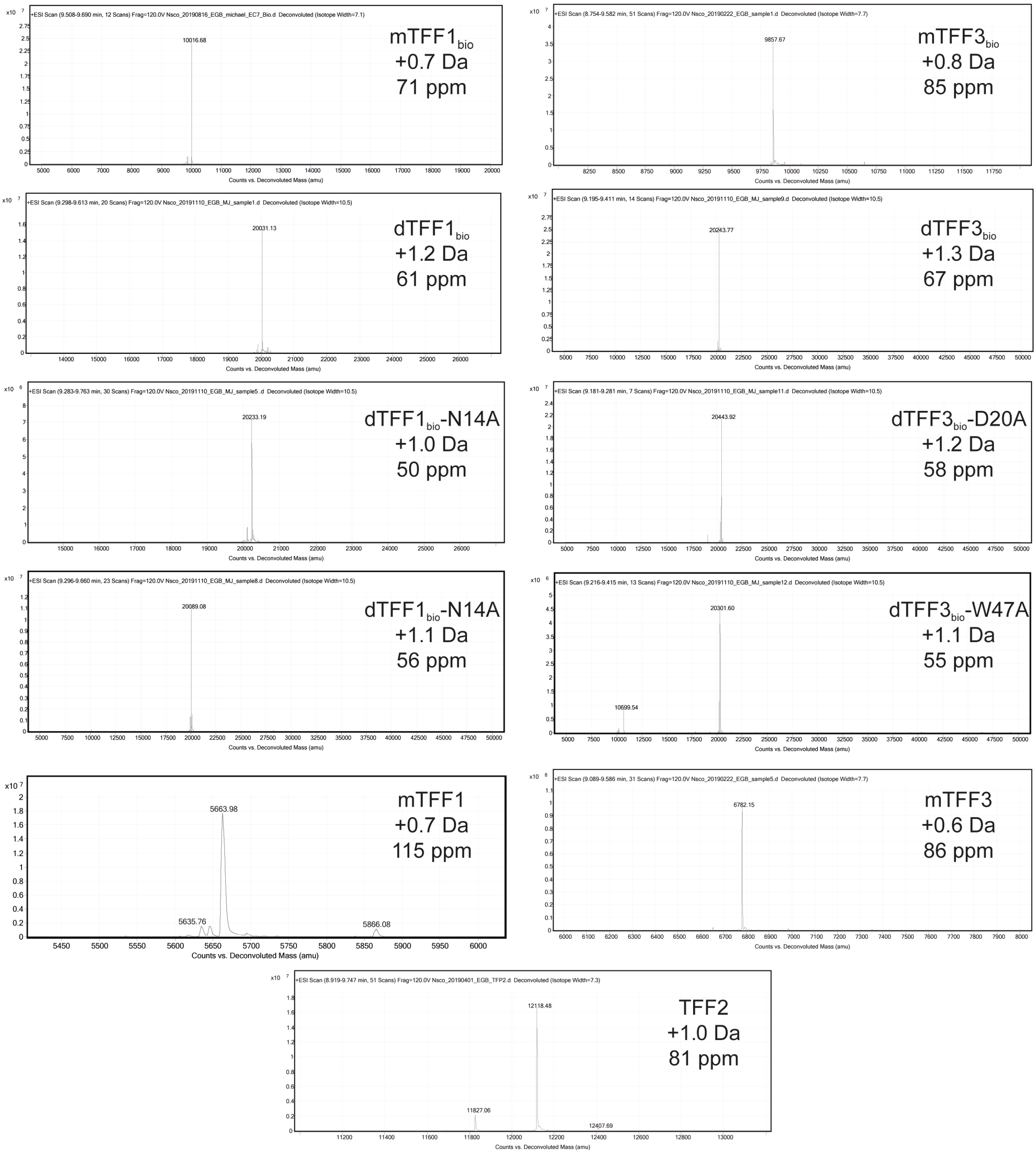

**Supplementary Figure 8.** Deconvoluted ESI-MS mass spectra of recombinant TFFs. The mass of reverse phase separated TFF are ±1.5 Da of the expected masses.
